# Distinct *Daphnia* spp. whole-body bacterial microbiota in two contrasting Mediterranean lakes

**DOI:** 10.64898/2026.03.31.714960

**Authors:** Vasileia Riga, Stefanos Katsoulis-Dimitriou, Eleni Nikouli, Maria Demetrzioglou, Evangelia Michaloudi, Konstantinos Kormas

**Author notes:** School of Veterinary Medicine, Aristotle University of Thessaloniki, Thessaloniki, Greece (Stefanos Katsoulis-Dimitriou).

## Abstract

The microbiota and microbiome associated with zooplankton remains rather understudied compared to other animal groups and/or taxa. The present study aimed at investigating the whole-body bacterial microbiota of *Daphnia* spp. in two contrasting Greek lakes, the shallow and hypertrophic Lake Koronia vs. the deep and mesotrophic Lake Vegoritida, including both egg-bearing and non-egg-bearing individuals. In both lakes, 2,060 bacterial operational taxonomic units (OTUs) were found, with 223 of them being conditionally rare (crOTUs) with low contribution even for the dominant phyla, with L. Vegoritida having more crOTUs than L. Koronia. The individuals microbiota had inconsiderable overlap with the surrounding water microbiota in both lakes. The two lakes showed significant differences in their *Daphnia* -associated microbiota. L. Koronia had richer OTUs and rather homogeneous bacterial communities, with higher occupancy. Overall, no significant differences in between the microbiota of egg-bearing and non-egg-bearing *Daphnia* individuals in both lakes. However, regarding the most important OTUs (miOTUs), the L. Koronia miOTUs were highly overlapped between the individuals with and without eggs, with only one missing from the individuals without eggs. In L. Vegoritida the individuals without eggs had only six miOTUs and while egg-bearing individuals had nine different ones; the two lakes had no shared miOTUs., considerable differences occurred.. A total of 27 miOTUs, was found and belonged to the Pseudomonadota, unclassified Bacteria, Cyanobacteria, Bacteroidota, Bacillota and Actinomycetota. Those miOTUs, where assignment to the genus level was possible, they were related to *Cyanobium*, *Mucilaginibacter, Flavobacterium* and *Staphylococcus*. This study showed that lake morphotype and ecological status can exert some impact on *Daphnia*-associated bacterial microbiota, with more pronounced effects on egg-bearing and non-egg-bearing individuals.

## INTRODUCTION

The water flea *Daphnia* is a genus of small planktonic crustaceans with over 100 species described and an extremely wide geographic distribution (Ebert 2022). The natural habitat of *Daphnia* varies from very small to large freshwater lakes and some genera are even found in saltwater lakes (Ebert 2022,). In aquatic food webs *Daphnia* plays a pivotal role as a primary consumer of phytoplankton and as a key food source for secondary consumers such as invertebrates and planktivorous fish (Miner et al. 2012). *Daphnia* species are often used as model organisms for ecology, evolution, physiology and ecotoxicology and as biomarkers for water quality (Reilly et al. 2023).

The well-being and physical condition of organisms are closely linked to the composition of their accompanying bacterial collectives. With the emergence and utilization of next generation DNA sequencing technologies (NGS), numerous studies about humans, animals and plants have proven the importance of microbiome on host health, nutrition and disease resistance (Ma et al. 2023). The gut microbiota specifically contributes to nutrient absorption, immune system regulation, metabolic regulation and disease prevention (Kim et al. 2025,).

The microbiota of plankton communities is shaped by a complex interplay of environmental, biological and ecological factors. Temperature (Jiang et al. 2019), nutrient availability (Velasquez et al. 2022), water mixing and flow (Wheeler et al. 2019), chemical and pollutant stressors (Xu et al. 2025), competition, predation and commensalism (Feng et al. 2025) are some of the factors that shape the microbiota of those communities. Host genotype and life-stage are primary determinants and diet plays a significant role in the gut microbiota assembly of zooplankton species (Xu et al.,. 2024). The microbiota of plankton communities is shaped by a complex interplay of environmental, biological and ecological factors like temperature, nutrient availability, diet, environmental stressors, competition and predation (e.g. Freese & Schink 2011, Macke et al. 2017,) and host genetics (Callens et al. 2020, Cooper & Cressler 2020, Gurung et al. 2024).

For *Daphnia*, the whole-body microbiota seems to be physiologically important as it has been found that when *Daphnia* individuals were raised without whole-body bacterial associates, they were smaller, less fecund and had higher mortality than those with microbiota (Sison-Mangus, 2015). Additionally, symbiotic bacteria in *D. magna* can contribute to increasing population size, the holobiont composition and host growth (Peerakietkhajorn et al. 2014, Callens et al. 2018). In *D. galeata*, the body microbiota can vary between genotypes (Rajarajan et al. 2023).

Lake depth causes changes in the configuration of these factors as deep lakes exhibit significant differences from shallow lakes in trophic structure and dynamics (Xue et al. 2025). Lake depth has a great influence on the plankton communities and ecosystem functions through light limitation, wind effect, nutrient dynamics, distribution patterns of plankton, and intensity of fish predation, thus it is expected to also cause changes in the microbiota assembly. Since genetic factors and life stage seem to play a role in shaping the *Daphnia* microbiota the presence or absence of eggs in the organism may also have an influence.

Most studies about *Daphnia*’s (mostly *D*. *magna*) microbiota are from lab populations and are focused on the gut (Akbar et al. 2022), although some studies about the whole-body microbiota exist (Callens et al. 2018, Cooper & Cressler 2020, Cooper et al. 2021,). Field studies with direct microbiota analysis from the natural environment samples are far more scarce (Hegg et al. 2021, Rajarajan et al. 2025). With the existing knowledge about the differences between laboratory *Daphnia* samples and field samples (Houwenhuyse et al. 2023) and the limited research on natural populations (Gurung et al. 2025), more information about the microbiota of wild *Daphnia* populations is needed.

The present study aimed at investigating the whole-body bacterial microbiota of *Daphnia* spp. natural populations in two contrasting Mediterranean lakes (shallow and hypertrophic vs. deep and mesotrophic). As the two lakes harbor different *Daphnia* species, we investigated the whole-body microbiota, in both egg-bearing and non-egg-bearing individuals, and its relation to the surrounding water microbiota rather than the host-species impact on shaping their whole-body microbiota.

## MATERIALS AND METHODS

### Sampling sites

Koronia is a shallow lake (max depth 1-3 m since 2010) located in northern Greece (23°04L-23°14L E, 40°7L-40°43LN, altitude: ca 75 m a.s.l., surface area 29 km^2^) (Figure S1). It is covered by the Directives 79/409/EEC and 92/43/EC, the RAMSAR Convention and is part of the National Wetland Park of Lakes Koronia-Volvi and the Macedonian Tempi (Michaloudi et al. 2012). The lake has dried up many times (2002, 2007, 2009, 2014), and bird and fish kills have occurred several times (1995, 2004, 2007, 2019) due to hypertrophic conditions and blooms of known toxic algal species (Genitsaris et al. 2009, Michaloudi et al. 2009).

Vegoritida (or Vegoritis) is a deep lake (max depth 52 m) with a surface area 46 km^2^ and belongs to the European Network of Protected Areas (NATURA 2000) (Fig. S1). It is located in northern Greece (40°47′ N, 513) ca. 115 km^2^ west of L. Koronia. Its trophic status has changed from oligotrophic at the beginning of the 1980s to meso-eutrophic at the end of the 2000s (Katsiapi et al. 2016), due to phytoplankton (cyanobacterial) blooms (Moustaka-Gouni & Nikolaidis 1990).

### Sample collection and processing

The samples analysed in the present study were collected in July 2019 from Lake Koronia (designated with K), and July/August 2019 from Lake Vegoritida (designated with V). Detailed information on the sampling procedure and in situ physicochemical parameter measurements is provided in Demertzioglou et al. (2022). In brief, zooplankton samples were collected from the deepest point (designated with D) as well as shallow littoral (designated with S) habitats, with vertical and horizontal hauls using plankton nets (50 and 100 μm mesh size). Upon collection, zooplankton samples were split into 50 ml Falcon tubes and were immediately (<1 min) flash-frozen in dry ice. During all samplings, in-situ measurements of Secchi disk water transparency and pH, using a HI-98100 Checker Plus pH Tester, were carried out (Table S1). Additional in-situ measurements included temperature and conductivity (μS/cm) using a YSI Pro 1030 instrument (YSI Inc., Ohio, USA). Water samples were collected from the water column using a 2-L Niskin sampler (50 cm height) at both lakes. Water samples were kept cool in the field and during transport to the laboratory and subsequently filtered through 0.20 μm membrane filters which were kept at-80^0^C until further analysis.

The zooplankton samples were transported to the laboratory and stored at −80°C. For the isolation of *Daphnia* individuals, each falcon tube was allowed to thaw on ice and individuals were randomly isolated under a stereoscope and a microscope. All procedures were conducted under sterile conditions to avoid sample contamination, while samples were kept on ice throughout the process. Female *Daphnia* individuals (*Daphnia magna* from L. Koronia and *Daphnia galeata* from L. Vegoritida) were selected from each falcon tube. At least 10 anatomically intact individuals from each sampling were chosen. Individuals with eggs (designated with +) and without eggs (designated with -) were separated. Each individual was placed in a 0.5 mL Eppendorf tube containing 100 μL of DNA/RNA Shield (Zymo Research, USA) and the samples were stored at 2-3^0^C until DNA extraction (<1 month).

Total whole-body DNA was extracted from six *Daphnia* individuals from each sample type and water filters by using the standard extraction protocol of the DNeasy PowerSoil Pro Kit (Qiagen, Germany). The only modification was the elution volume which was set to 40 μl. The V3–V4 regions of the bacterial 16S rRNA genes were amplified from the extracted DNA with the primer pair S-D-Bact-0341-b-S-17 and S-D-Bact-115 0785-a-A-21 (Klindworth et al. 2012). Sequencing of the amplicons was performed on a MiSeq Illumina instrument (2×300 bp) at the MRDNA Ltd. (Shallowater, TX, USA) sequencing facilities. The raw DNA sequences from this study have been submitted to the Sequence Read Archive (https://www.ncbi.nlm.nih.gov/sra/) in the BioProject PRJNA1237258 (BioSample SAMN47415465). The standard operating procedure of MOTHUR software (v.1.48.0) (Schloss et al. 2009, Schloss et al. 2011) was used for processing all the raw 16S rRNA sequence reads. Sequences assigned as mitochondria and chloroplasts, and single singletons were excluded from further analyses. The operational taxonomic units (OTUs) were determined at 97% cutoff similarity level and were classified with the SILVA database release 138.2 (Quast et al. 2013, Yilmaz et al. 2014). The final OTUs table was normalized to 26,726 sequence reads per sample. The Nucleotide BLAST (http://blast.ncbi.nlm.nih.gov) tool was used for identifying the closest relatives of the resulting OTUs.

The PAleontological STudies (PAST) software (Hammer et al. 2001) was used for calculating diversity indices and testing of the statistically significant differences across all samples by applying non-metric multidimensional scaling (nMDS) based on the unweighted pair group method with the arithmetic mean Bray–Curtis similarity. Additionally, the Bray-Curtis similarity permutational multivariate analysis of variance (PERMANOVA) with 9999 permutations, between the four sampling treatments, was applied.

Based on the distribution of each OTU’s abundance in the whole dataset, each OTU was assigned as conditionally rare (crOTUs) when its coefficient of bimodality b was ≥0.9 and relative abundance ≥0.01%. The coefficient of bimodality b was calculated according to the following equation, where x is the abundance of each OTU in sample i,

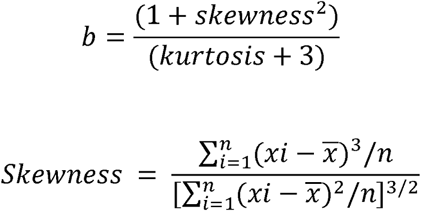

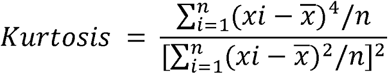

## RESULTS

The 1,385,973 raw sequences were assigned to 2,060 bacterial OTUs across all water at *Daphnia* individuals and water samples. The highest OTUs richness (211-235) was found in L. Koronia compared to L. Vegoritida (126-134) (Table 1). Lake Koronia had also the highest alpha diversity indices of Simpson 1-D (0.88-0.92) and Shannon (H) (3.06-3.26) contrary to L. Vegoritida (0.60-0.90 and 1.91-3.16, respectively). Evenness was comparable in both lakes, 0.09-0.13 and 0.08-0.19 in L. Koronia and L. Vegoritida, respectively. Lake Vegoritida harboured highest percentages of crOTUs (8.9-15.2%) compared to L. Koronia (6.6-7.8%). All water samples had much higher OTUs richness, alpha diversity indices and percentage of crOTUs, with comparable evenness values. No statistically significant differences were found between *Daphnia* individuals with and without eggs in each lake (Table 2).

**Table 1.** Diversity indices (average ± standard deviation) of the Daphnia associated bacterial operational taxonomic units (OTUs).

|  | Koronia |  |  |  | Vegoritida |  |  |  | Koronia |  | Vegoritida |  |
| --- | --- | --- | --- | --- | --- | --- | --- | --- | --- | --- | --- | --- |
|  | Eggs + |  | Eggs - |  | Eggs + |  | Eggs - |  | Water |  | Water |  |
|  | Deep | Shallow | Deep | Shallow | Deep | Shallow | Deep | Shallow | Deep | Shallow | Deep | Shallow |
| OTUs | 211 | 223 | 235 | 226 | 126 | 134 | 130 | 132 | 471 | 480 | 450 | 502 |
| (S) | $\pm 29$ | $\pm 49$ | $\pm 32$ | $\pm 52$ | $\pm 21$ | $\pm 16$ | $\pm 31$ | $\pm 23$ | | | | |
| Simpson 1-D | 0.92 | 0.88 | 0.89 | 0.89 | 0.60 | 0.90 | 0.79 | 0.71 | 0.96 | 0.94 | 0.95 | 0.96 |
| | $\pm 0.01$ | $\pm 0.09$ | $\pm 0.02$ | $\pm 0.03$ | $\pm 0.37$ | $\pm 0.19$ | $\pm 0.21$ | $\pm 0.17$ | | | | |
| Shannon | 3.26 | 3.08 | 3.06 | 3.10 | 1.91 | 3.16 | 2.47 | 2.17 | 4.07 | 3.74 | 4.26 | 4.02 |
| (H) | $\pm 0.14$ | $\pm 0.62$ | $\pm 0.15$ | $\pm 0.27$ | $\pm 1.19$ | $\pm 0.52$ | $\pm 0.78$ | $\pm 0.78$ | | | | |
| Evenness | 0.13 | 0.11 | 0.09 | 0.11 | 0.08 | 0.19 | 0.11 | 0.08 | 0.12 | 0.09 | 0.16 | 0.11 |
| (eH/S) | $\pm 0.03$ | $\pm 0.06$ | $\pm 0.02$ | $\pm 0.05$ | $\pm 0.07$ | $\pm 0.07$ | $\pm 0.06$ | $\pm 0.06$ | | | | |
| Conditionally<br>rare OTUs | 7.8 | 7.0 | 7.0 | 6.6 | 8.9 | 11.5 | 11.4 | 15.2 | 11.4 | 9.6 | 18.0 | 14.8 |
| (%) | $\pm 0.9$ | $\pm 0.8$ | $\pm 1.9$ | $\pm 0.8$ | $\pm 1.3$ | $\pm 2.1$ | $\pm 2.6$ | $\pm 2.6$ | | | | |

**Table 2.** Results of permutational analysis of variance between the Daphnia microbiota. PERMANOVA. K: Lake Koronia, V: Lake Vegoritida, +/-: with eggs/without eggs, D: deep, S: shallow water. Red indicates p < 0.05. Upper-right half: F values, lower-left half: p values.

|  | *K+D* | *K+S* | *K-D* | *K-S* | *V+D* | *V+S* | *V-D* | *V-S* |
| --- | --- | --- | --- | --- | --- | --- | --- | --- |
| **K+D** | - | 0.828 | 0.406 | 1.211 | 3.465 | 6.958 | 4.264 | 10.850 |
| **K+S** | 0.733 | - | 0.912 | 0.896 | 2.974 | 4.363 | 3.312 | 7.882 |
| **K-D** | 0.819 | 0.500 | - | 1.020 | 3.673 | 7.386 | 4.659 | 10.900 |
| **K-S** | 0.322 | 0.578 | 0.398 | - | 3.643 | 7.379 | 4.313 | 10.860 |
| **V+D** | 0.003 | 0.002 | 0.003 | 0.002 | - | 2.090 | 0.851 | 3.439 |
| **V+S** | 0.003 | 0.002 | 0.002 | 0.002 | 0.105 | - | 2.940 | 2.650 |
| **V-D** | 0.003 | 0.003 | 0.003 | 0.002 | 0.233 | 0.002 | - | 4.985 |
| **V-S** | 0.002 | 0.002 | 0.002 | 0.003 | 0.015 | 0.097 | 0.002 | - |

Based on the nMDS topology, there was a distinct separation of both lakes *Daphnia*’s microbiota and their respective water samples (Figure 1). This was also confirmed by the statistically significant differences between the two lakes (Table 2). The L. Koronia *Daphnia*’s microbiota was considerably more overlapped between the different sample types. Lake Koronia had OTUs with high occupancy compared to L. Vegoritida (Figure 2).

**Figure 1.**
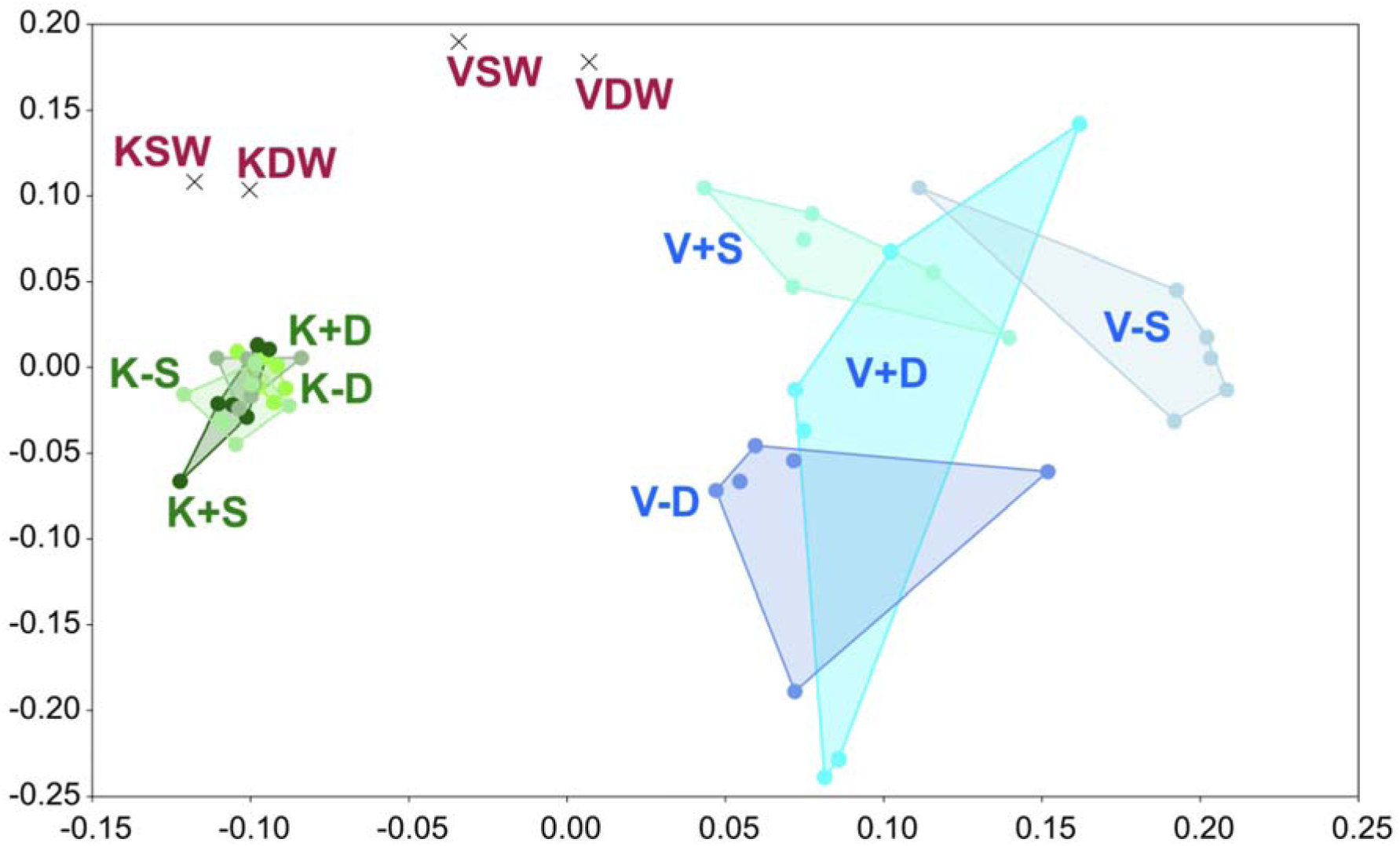
Non-metric multidimension scaling of the water (W), *Daphnia* microbiota from shallow (S) and deep (D) waters of lakes Koronia (K) and Vegoritida (V), Greece. +: with eggs, -: without eggs.

**Figure 2.**
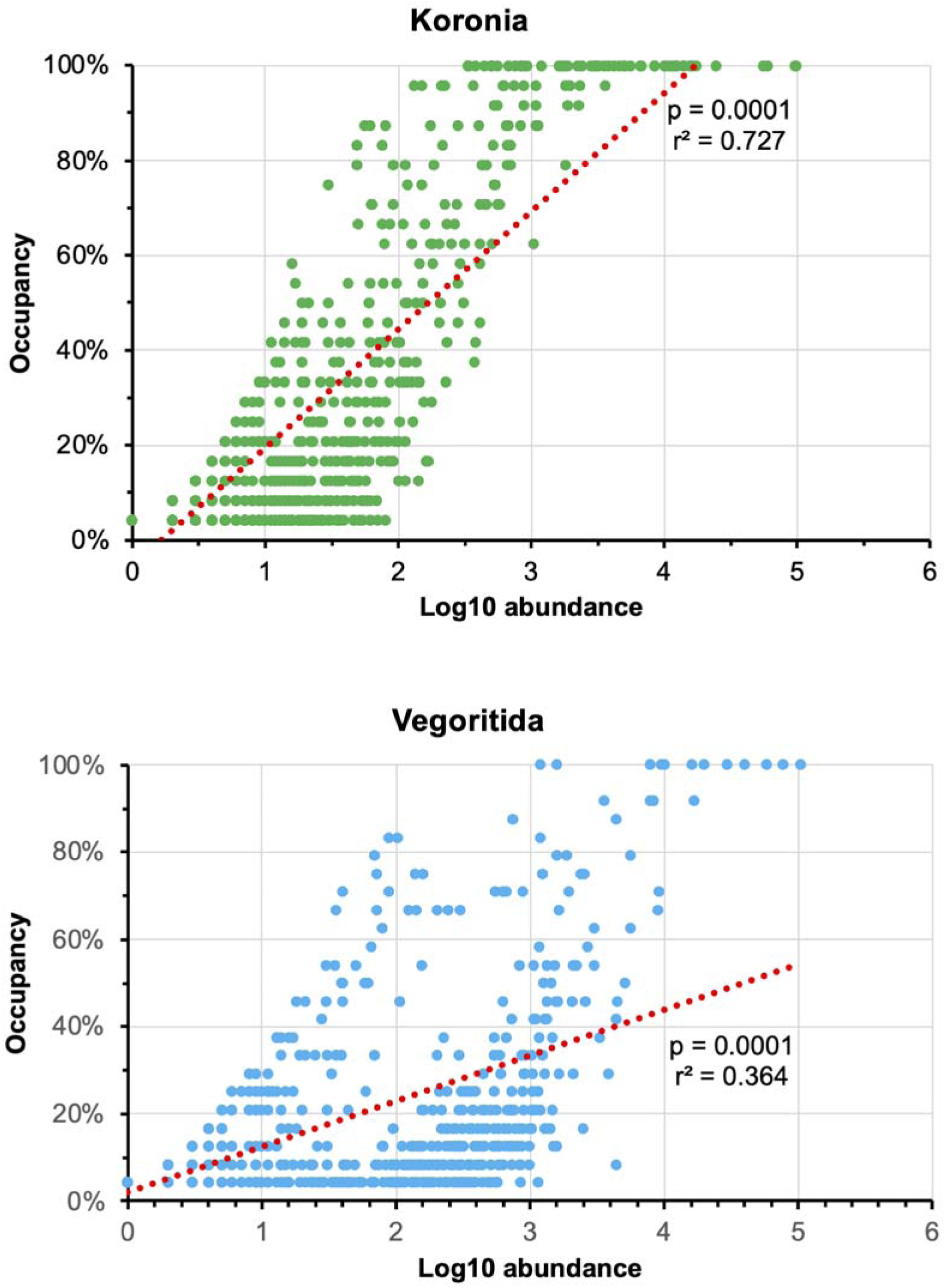
Occupancy of the *Daphnia* spp. whole-body bacterial microbiota found in two Mediterranean lakes, Greece.

A total of 30 bacterial taxa at the phylum level was found to be associated with *Daphnia* in both lakes (Figure 3). In L. Koronia, the Pseudomonadota dominated by 35.9-45.0%, followed by unclassified Bacteria (15.9-19.4%), Cyanobacteriota (6.3-15.7%) and Bacteroidota (7.2-10.0%). In L. Vegoritida the Pseudomonadota (27.9-54.0%), Bacteroidota (21.8-28.1%), Bacillota (7.6-19.7%) and Actinomycetota (5.7-16.8%) dominated.

**Figure 3.**
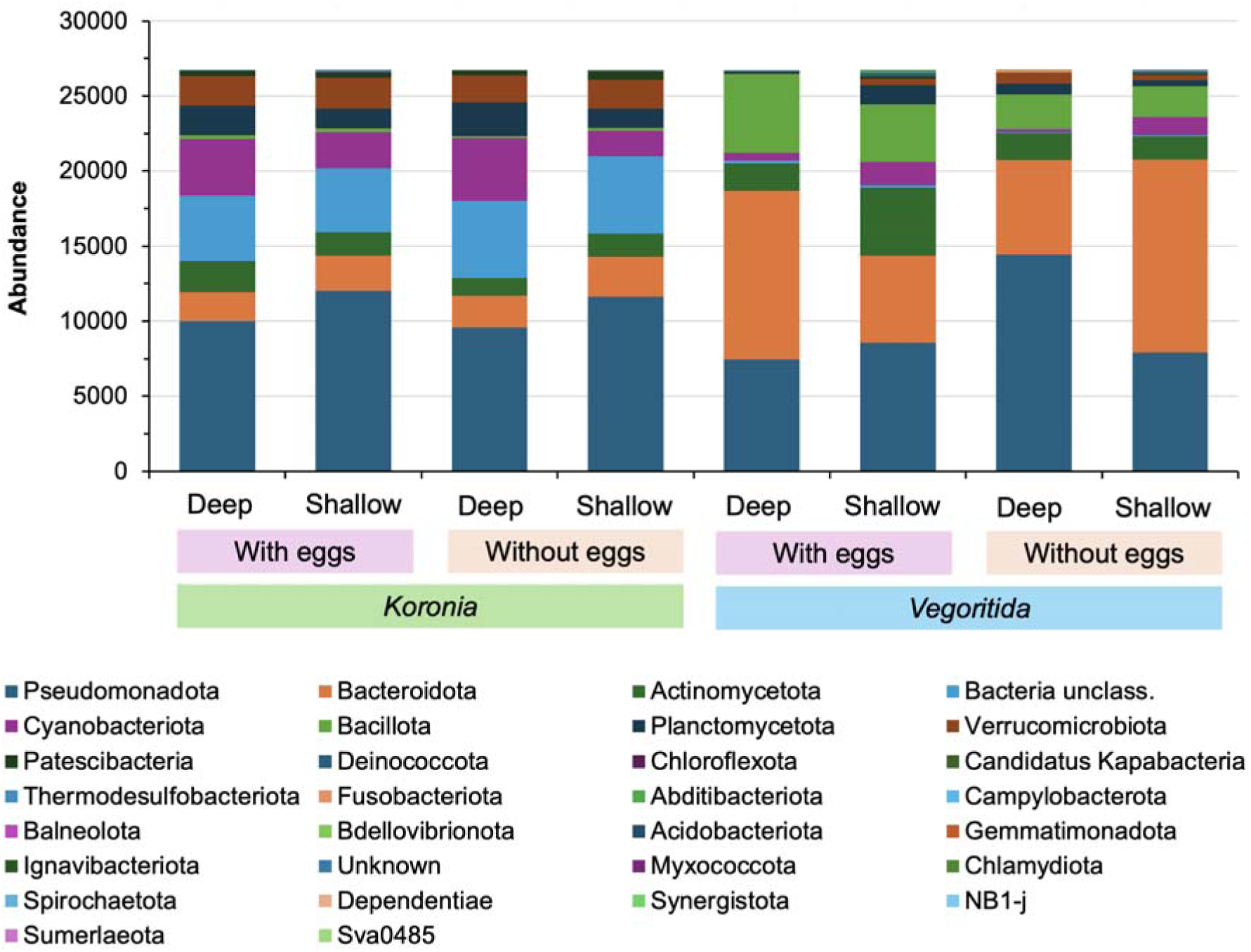
Bacterial phyla found in the *Daphnia* spp. whole-body microbiota.

The OTUs with ≥1% relative abundance per sample, in all L. Koronia individuals (Figure 4) were affiliated - in descending abundance order- to the Paracocacceae family (OTU0010, −4, −15, 50; 17.1-25.3% relative abundance), unknown bacterial families (OTU0003, −30; 14.1-18.1%), Comamonadaceae family (OTU0005, −20, −151; 10.4-11.7%) and the genus *Cyanobium* (OTU0006, −9; 4.4-14.9%). In L. Vegoritida, different OTUs dominated the various *Daphnia* individuals (Figure 5). *Mucilaginibacter*-related OTU0001 (15.4-32.1%) dominated in V+D and it was the third most abundant in V-D, followed by *Staphylococcus* (OTU0008; 2.6-10.9%) and unknown Burkholderiales (OTU0007, 12; 0.9-18.4%). The latter order was dominant in V-D followed by unknown Comamonadaceae (OTU0005, −20, −151; 1.2-16.2%). Samples V+S and V-S were dominated by *Flavobacterium* OTUs (OTU0002, −86, 97, 190; 5.1-41.8%), followed by *Staphylococcus* (OTU0008; 2.6-10.9%) and unknown Burkholderiales (OTU0007, 12; 0.9-18.4%).

**Figure 4.**
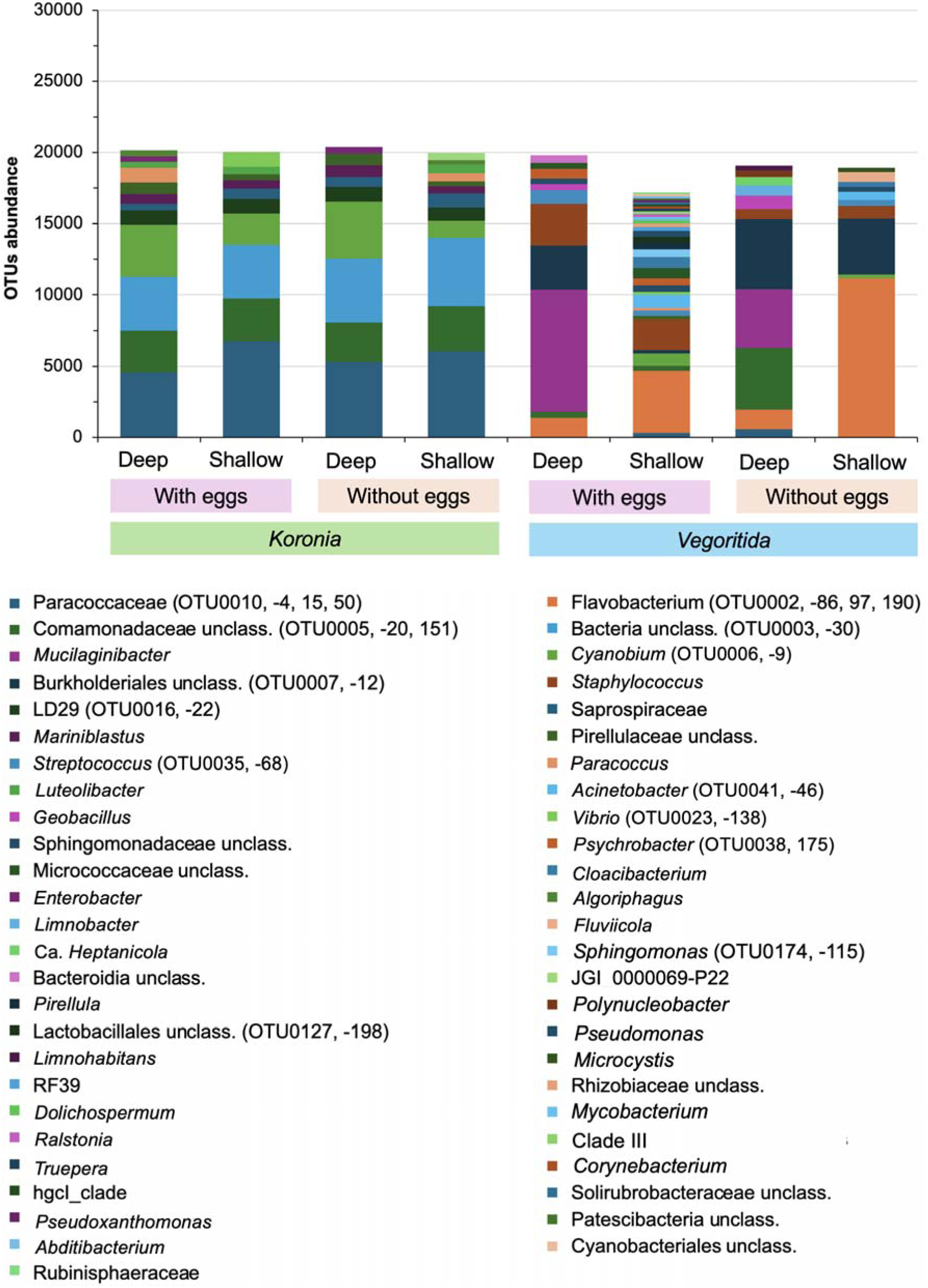
Genus-level of the bacterial most important operational taxonomic units (OTUs) found in the *Daphnia* whole-body microbiota with ≥1% relative abundance.

**Figure 5.**
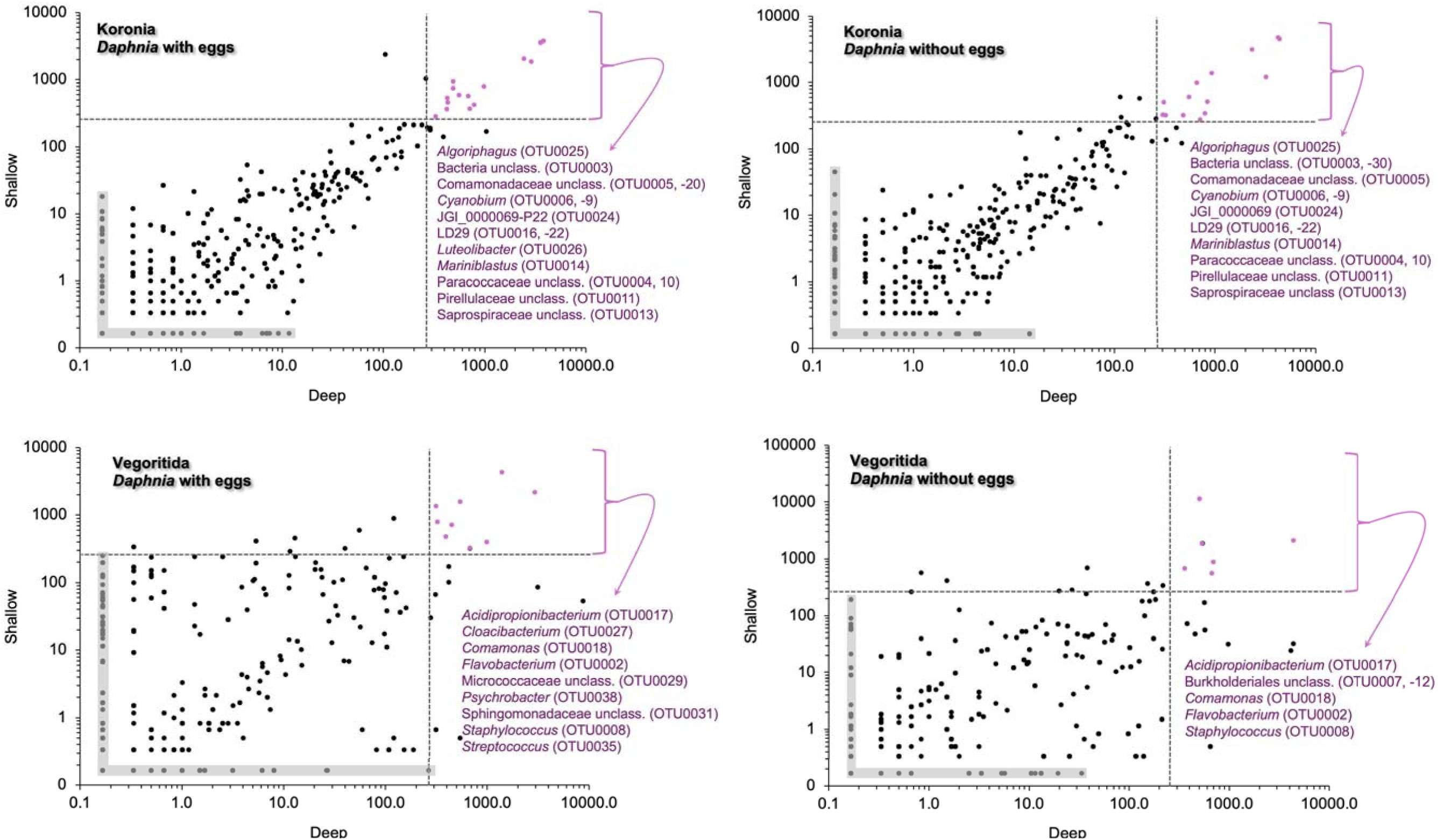
Most important bacterial operational taxonomic units (OTU) (in purple) at the genus level found in the *Daphnia magna* whole-body crobiota. Dashed lines indicate ≥1% relative abundance. Shaded areas contain OTUs with no abundance in one of the two habitats.

In total, 223 OTUs were characterized as crOTUs but their contribution was very low even for the dominant (≥1% relative abundance) phyla, ranging between 0.05% (one OTU in all Chloroflexota OTUs) and 4.0% (83 OTUs in all Pseudomonadota OTUs) (Table S2). Twenty-nine crOTUs belonged to the most abundant (>1% relative abundance) in the *Daphnia* individuals (Table S3). In this group of 29 OTUs, OTU0015 (Paracocacceae) was included among the L. Koronia miOTUs, while OTU0001 (*Mucilaginibacter*), −86, −97 and −190 (*Flavobacterium*) were included in the L. Vegoritida miOTUs. In the present study, we define as “most important OTUs” (miOTUs) these OTUs which had concomitantly ≥1% relative abundance in the individuals from both the shallow and deep habitats. A total of 27 miOTUs were found belonging to 21 bacterial taxa at the genus, family, order or unclassified level (Figure 5). The miOTUs of L. Koronia *Daphnia* individuals were highly overlapped between the individuals with and without eggs, with only one (the *Luteolibacter*-related OTU0026) missing from the individuals without eggs. Similarly, in L. Vegoritida, the individuals without eggs had only six miOTUs while egg-bearing individuals had nine. The two lakes had no shared miOTUs.

In L. Koronia, 15.6-15.8% of all OTUs where shared between the water and *Daphnia*-associated bacterial communities, while the respective range for L. Vegoritida was 9.6-10.8% (Figure S2). Based on the Sorensen-Dice index, the whole-body microbiota of non-egg-bearing *Daphnia* from the shallow L. Vegoritida, showed the lowest similarity with all individuals from L. Koronia and the deep L. Vegoritida. The rest of the individuals had medium to high similarity between them (Table 3).

**Table 3.** Sorensen-Dice index (scaled between 0 and 1) of the whole-body bacterial operational taxonomic units (OTUs) associated with *Daphnia* spp. individuals with (+) and without (-) eggs from the shallow (S) and deep (D) parts of lakes Koronia (K) and Vegoritida (V), Greece. Green: Low similarity (0.20-0.39), pink: medium similarity (0.40-0.59), orange: high similarity (0.60-0.79) between bacterial communities.

|  | *K+D* | *K+S* | *K-D* | *K-S* | *V+D* | *V+S* | *V-D* | *V-S* |
| --- | --- | --- | --- | --- | --- | --- | --- | --- |
| *K+D* | - |  |  |  |  |  |  |  |
| *K+S* | 0.57 | - |  |  |  |  |  |  |
| *K-D* | 0.62 | 0.59 | - |  |  |  |  |  |
| *K-S* | 0.56 | 0.63 | 0.59 | - |  |  |  |  |
| *V+D* | 0.65 | 0.60 | 0.48 | 0.45 | - |  |  |  |
| *V+S* | 0.64 | 0.59 | 0.47 | 0.44 | 0.61 | - |  |  |
| *V-D* | 0.43 | 0.52 | 0.55 | 0.40 | 0.56 | 0.54 | - |  |
| *V-S* | 0.29 | 0.27 | 0.33 | 0.27 | 0.37 | 0.40 | 0.47 | - |

## DISCUSSION

In this study, the whole-body bacterial microbiota of two distinct populations of *Daphnia* spp. was characterized in two different Mediterranean lakes in Greece. L. Koronia had higher OTUs richness and alpha diversity indices than Lake Vegoritida, while L. Vegoritida contained more crOTUs. Evenness values were low and comparable in the two lakes. The higher OTUs richness and alpha diversity indices of *Daphnia* individuals from L. Koronia can be explained due to the increased availability of growth-limiting resources in a hyper-eutrophic shallow ecosystem (Xie et al. 2024). Eutrophic lakes have been previously found to have higher microbial diversity than oligotrophic ones (Sauer et al. 2022). However, studies have also shown contrasting results. A freshwater eutrophic system does not always support higher alpha diversity than an oligotrophic one, but changes in the structure of microbial communities (beta diversity) always exist (Yanez-Montalvo et al. 2022, Kubera et al. 2025). Sometimes, a eutrophic environment can have decreased alpha diversity, especially when cyanobacterial blooms exist that favour only certain bacterial species (Huang et al. 2022). However, the host possibly plays a role at filtering that diversity and choosing certain microbial species as its associates (Wall et al. 2025).

Higher microbial diversity suggests better host health, although that may not be the case in a eutrophic lake, where elevated diversity could also reflect environmental stressors (Zhang et al. 2025). In contrast, in an oligotrophic lake, the microbiome may be more stable with more beneficial taxa for the host (Jackrel et al. 2023). However, a greater number of crOTUs were observed in Lake Vegoritida. These crOTUs are microbial taxa that are present at very low abundances but have the potential to become dominant under certain environmental or biological conditions. Vegoritida is a mesotrophic lake, but it experiences sudden events such as cyanobacterial blooms, probably when nutrients from agricultural activities or very high temperatures exist in the area (Zervou et al. 2021, Tsitsis et al. 2023). Conditions change abruptly from oligotrophic to eutrophic, resulting in crOTUs that reside in *Daphnia* hosts and can become abundant. Frequent and prolonged harmful blooms are also happening in L. Koronia, but organisms such as *Daphnia* and the plankton communities are possibly already adapted to this hyper-eutrophic environment, thus, the microbiota becomes more stable and involves less crOTUs. The microbiota of L. Vegoritida organisms can possibly respond more dynamically to environmental changes and reorganize in response to stressful conditions or nutrient changes.

There was a clear and statistically significant separation, based on nMDS and PERMANOVA, between the two lakes Daphnia-associated microbiota and their water samples. Also, nMDS showed the high overlap of the microbiota in all *Daphnia* individuals from L. Koronia and, conversely, the more dispersed microbiota in individuals from L. Vegoritida. The separation of water microbiota and the microbiota of *Daphnia* individuals contrasts with the filter feeding behavior of *Daphnia* and with the whole-body microbiota that is in direct contact with water, suggesting that the Daphnia associated bacteria are gained from other routes as well. Similar studies have found that the water microbiome differed from the gut microbiome of *Daphnia* (Freese & Schink, 2011, Hegg et al. 2021), as well as other species that exhibit filter feeding behaviour such as sponges (Taylor et al. 2012), bivalves (Alvanou et al. 2025), oysters (Arfken et al. 2021) and ascidians (Galia-Camps et al. 2023). This proves that the microbiota is not selected from the environment and water, but rather that bacteria useful for the organism’s functions or those capable of surviving in the conditions of the *Daphnia*’s intestine and body are selected. Therefore, some OTUs, which are found in small concentrations in the water, may pass into *Daphnia* and dominate its microbiota. All miOTUs were also present in the water microbiota in much smaller concentrations than in the *Daphnia* microbiota. Another possible explanation is that the animals may be colonized by water bacteria that were not present in high abundances in the water samples but existed in previous days/weeks.

In terms of bacterial phyla, there were several similarities between the two *Daphnia* species in the two different lakes. The phylum Pseudomonadota (formerly Proteobacteria) is the most common bacterial phylum in *Daphnia* datasets, specifically the Betaproteobacteria and Gammaproteobacteria (Peerakietkhajorn et al. 2014, Callens et al. 2018, Rajarajan et al. 2023). Bacteroidota and Actinomycetota are also dominant phyla and appear consistently in high relative abundances in *Daphnia* datasets (Cooper & Cressler, 2020, Varg et al. 2022). Cyanobacteriota and Bacillota are also common, although in lower relative abundances (Macke et al. 2017, Mushegian et al. 2019, Hegg et al. 2021).

The differences in the bacterial composition of the *Daphnia* populations of the two lakes, which were evident in the alpha and beta diversity indices, are better understood when considering lower taxa, i.e. genus and family levels. More specifically, the individuals in L. Koronia showed minimal differences between them in terms of the genera and families that appeared, while in L. Vegoritida, depth appears to play an important role in shaping bacterial genera and families. The minimal differences in L. Koronia reinforce the observation about the importance of depth, as it is a shallow lake with small differences in depth across its surface (Mitraki et al. 2004), in contrast to Vegoritida, which is a deep ecosystem (Skoulikidis et al. 2008). Another important observation is the minimal effect that eggs had on the formation of the animal’s microbiota in both lakes, which contrasts with the knowledge about the role of life stage, host, and reproduction in *Daphnia* (Mushegian et al., 2019). As can be seen, in populations in their natural environment, the biological factor of the presence of eggs has less influence than the environment in which *Daphnia* individuals develop.

Lake Koronia was dominated by unclassified Bacteria of the Paracoccaceae family, which could not be affiliated to any known taxa of this family. BLAST results gave identities between 77.3% and 89.2% with Paracoccaceae sequences in GenBank. This family contains taxa which thrive in a wide range of environments including freshwater, marine systems, soil, sediments, and host-associated microbiomes (Hollesteiner et al. 2023). Alphaproteobacteria are not the most abundant class reported in *Daphnia* studies and Paracocacceae seems to be a rare family. Paracoccaceae involves species from polluted or nutrient rich environments with versatile metabolic attributes that contribute to nitrogen cycling or organic matter decomposition (Olaya-Abril et al. 2018,). In eutrophic systems such as L. Koronia nutrient levels and phytoplankton-derived substrates are in high concentrations making the environment favorable for Paracoccaceae family members. The Comamonadaceae family contains aerobic or facultatively anaerobic heterotrophs that are frequently among the dominant bacterial taxa in freshwater lake plankton (Willems, 2013). It is an important family in *Daphnia* microbiota and it contains one of the most important *Daphnia*-associated microbiota, the genus *Limnohabitans,* which is represented by OTU0005 in this study as consistently dominant in *Daphnia* microbiota studies. This genus is considered a beneficial symbiont to *Daphnia*, enhancing growth, reproduction, immune system and resilience to environmental stress (Peerakietkhajorn et al. 2016, Cooper & Cressler, 2020). The other two Comamonadaceae OTUs (−20, −151) were distantly affiliated with known genera or species of this family (BLAST identities ≤94%). *Cyanobium* is a genus of freshwater cyanobacteria that perform photosynthesis and thrive in hyper-eutrophic, warm and high-light environments (Kwon et al. 2020). High nutrient concentrations such as nitrogen and phosphorus can lead to cyanobacterial blooms and high abundance of the *Cyanobium* genus (Kramer et al. 2024). *Cyanobium* can be direct food for *Daphnia* and reside in the gut and it also constitutes a biomarker of eutrophic environments, consistent with the trophic status of L. Koronia.

In L. Vegoritida, OTU0001 which belonged to the genus *Mucilaginibacter,* was the most abundant in the animals from the deep part of the lake, followed by OTU0007 (Burkholderiales unclassified, but had 99.1% sequence similarity to an OTU from lake water with GenBank accession number MZ246148) and OTU0008 (*Staphylococcus*). In our case, the *Limnohabitans*-related OTU0005 was the second most abundant in the individuals without eggs. The *Daphnia* individuals from the shallow samples were dominated by *Flavobacterium* OTUs, although *Staphylococcus* and Burkholderiales also had a strong presence. *Mucilaginibacter* species have been isolated from various environments including soil, water and plants (Hirsch et al. 2024, Li et al. 2025). Several species of this genus are known to degrade polysaccharides and complex carbon (Kumar et al. 2025). Consequently, their strong presence in the animals from the deep habitat and not in the shallow one, can be attributed to the higher concentrations of organic carbon that sometimes exist in the deeper parts of a lake (Li et al. 2021). Burkholderiales are commonly found in *Daphnia* datasets and are linked with host growth and nutrient assimilation (Qi et al. 2009, Rajarajan et al. 2025). *Staphylococcus* is another not common genus in *Daphnia* microbiota studies, although it is found in low abundances (Cooper & Cressler, 2020). It contains pathogenic species for humans, but it commonly occurs in aquatic environments (Sánchez et al. 2017). It is found in marine copepods (Feng et al. 2023) and it is considered a beneficial genus in fish gut (Katsoulis-Dimitriou et al. 2024). Based on this information, *Staphylococcus* seems to be a beneficial genus for *Daphnia* microbiota, possibly occurring in the gut bacterial communities. *Flavobacterium* OTUs were the most abundant in the shallow samples. *Flavobacterium* genus is identified as a core component of the *Daphnia* microbiome (Cooper & Cressler, 2020). It is linked with increased tolerance to pesticides in *Daphnia* hosts and the degradation of cyanobacterial toxins (Macke et al. 2017, Janseens et al. 2022). *Flavobacterium*, also contains species with the ability to digest phytoplankton cell wall polysaccharides, thus it may contribute to nutrient acquisition for *Daphnia* hosts (Gavriilidou et al. 2020).

Several OTUs were classified as miOTUs in the two lakes. In L. Koronia individuals, *Mariniblastus*, *Algoriphagus*, *Cyanobium*, LD29, Paracoccaceae, Patescibacteria, Saprospiraceae, Pirellulaceae, Comamonadaceae (*Limnohabitans*) and unclassified Bacteria were present in high abundances in all habitats and hosts, while *Luteolibacter* was abundant concomitantly in individuals with eggs form the deep and shallow parts of the lake. *Mariniblastus* is a genus that has been found only in marine environments attached to macroalgae biofilms (Lage et al. 2017). Interestingly, it is the first time reported in a freshwater ecosystem and in *Daphnia* microbiota, thus its function is unknown. It is speculated that increased conductivity of L. Koronia at the time of sampling (6.2 – 12.2 mS cm^-1^; Michaloudi unpubl. data) caused by abrupt reduction of its water level, could favor the abundance of marine origin taxa from the nearby Thermaikos Gulf. *Algoriphagus*, LD29, *Cyanobium* and *Luteolibacter* are all genera that occur in freshwater environments and likely reflect dietary intake or environmental acquisition in *Daphnia* microbiota (Han et al. 2017, Szabó et al. 2020, Farkas et al. 2022). OTU0010 from the Paracocacceae family was assigned to *Cypionkella collinsensis*, a freshwater aerobic bacterium (Hördt et al. 2020) with minimal information about its potential role in zooplankton microbiome. The closest relative of the other Paracoccaceae bacterium (OTU0004) was assigned to *Frigidibacter*, a genus with species isolated from freshwater environments (Li & Zhou, 2015) and from oil contaminated water (Zhang et al. 2020). Patescibacteria and Saprospiraceae are inhabiting freshwater, brackish and saline environments and are associated with microalgae and activated sludge (Vettorazzo et al. 2024, Mcllroy & Nielsen 2014). Saprospiraceae are also found in various hosts, including marine sponges and algae (Hosoya et al. 2006). Pirellulaceae are more common in marine habitats, but they also contain some freshwater species (Kulichevskaya et al. 2022). They have been found in hosts such as corals, marine sponges and ascidians (Goldsmith et al. 2018, Matos & Antunes, 2021, Vitorino et al. 2022). Without further identification it is difficult to know the exact role of these OTUs on *Daphnia* microbiota, but since they have been found in aquatic environments and associated with various hosts they possibly have a beneficial role for Daphnia individuals.

In Lake Vegoritida *Comamonas*, *Acidipropionibacterium*, *Staphylococcus* and *Flavobacterium* were abundant in all individuals, while Micrococcaceae (*Kocuria* sp.), *Cloacibacterium*, *Psychrobacter*, Sphingomonadaceae (*Sphingopyxis* sp.) and *Streptococcus* were abundant concomitantly in individuals with eggs from the deep and shallow habitats. *Streptococcus* has been associated with the gut habitat of freshwater crustacean zooplankton (Li et al. 2026). Burkholderiales was abundant concomitantly in Deep and shallow samples and in the samples without eggs. *Comamonas* spp. have been isolated from a broad variety of environments, including water, aircraft water, soil, plants, and animals and have been investigated for their ability to degrade xenobiotic pollutants and for heavy metal detoxification (Ryan et al. 2022). *Acidipropionibacterium* are bacteria that produce propionic acid via fermentation, and they have also been found to produce vitamin B12 (Miyamoto et al. 2025). Propionate is a short chain fatty acid, a gut bacterial metabolite that can act as energy source for the host and has several beneficial effects such as improvement of energy metabolism and reduction of cholesterol (Rahman et al. 2023). From the available information *Comamonas*, *Acidipropionibacterium*, *Staphylococcus* and *Flavobacterium* all possibly play important beneficial roles for *Daphnia* hosts.

Micrococcaceae is a family found in the *Daphnia* microbiota in small percentages (Choi et al. 2024). *Kocuria* contain species that colonize the skin in mammals, and some are human pathogens (Ziogou et al. 2023). They are found in freshwater and marine environments and *Kocuria rhizophila* is known to cause disease in salmonid species (Pekala-Safinska et al. 2017). *Cloacibacterium* is a genus containing species isolated from wastewater or lake sediment, that can produce extracellular polymeric substances (Klai, 2016). *Cloacibacterium haliotis* has been isolated from the intestine of abalone, showing the possible role of this genus in the gut microbiota of organisms (Hyun et al. 2014). Members of the genus *Psychrobacter* thrive in marine cold environments or in cold freshwater lakes (Rodrigues et al. 2009). They are also found in the gut microbiota of freshwater and marine fish (Makled et al. 2017, Diwan et al. 2022). Sphingomonadaceae contains bacteria such as *Sphingomonas* that produce sphingolipids that are known to modulate the mammalian immune system (Heaver et al. 2018). *Sphnigopyxis* sp. inhabit a variety of environments from agricultural soil to marine and freshwater and harbour bio-degradative capabilities for various environmental contaminants (Sharma et al. 2020). *Sphingomonas* also alleviates silver nanoparticle toxicity in *D. magna* (Ouwehand et al. 2025). *Streptococcus* is a genus that has been reported again in *Daphnia* microbiota in low abundances (Bulteel et al. 2021).

## ACKNOWLEDGEMENTS

This work was financially supported by the English MSc Program “Host-Microbe Interactions” of the University of Thessaly, Greece.

