## Supplementary material for "Distinct *Daphnia* spp. whole-body bacterial microbiota in two contrasting Mediterranean lakes"

**Table S1**. Physical and chemical parameters in lakes Koronia and Vegoritida, Greece in 2019. d.l.: detection limit, TN: total nitrogen, TON: total organic nitrogen, SRP: soluble reactive phosphorus, TP: total phosphorus.

| | **Koronia (K)** | | **Vegoritida (V)** | | | |
|---|---|---|---|---|---|---|
| | July | | July | | August | |
| | Deep | Shallow | Deep | Shallow | Deep | Shallow |
| pH | 8.77 | 8.83 | 9.06 | 8.72 | 9.48 | 9.49 |
| Temperature (°C) | 27.8 | 28.2 | 24.2 | 26.6 | 25.3 | 26.8 |
| Conductivity (μS $cm^{-1}$) | 8800 | 8816 | 785 | 749 | 799 | 795 |
| Salinity (ppt) | 4.6 | 4.6 | < d.l. | | < d.l. | |
| Max depth (m) | 1 | 1 | 43 | | 40 | |
| Transparency (Secchi m) | 0.3 | 0.3 | 3.8 | | 2.9 | |
| N-$NO_2^-$ (mg N $^{-1}$) | 0.017 | | 0.003 | | 0.003 | |
| N-$NO_3^-$ (mg N $L^{-1}$) | 0.21 | | 0.159 | | 0.224 | |
| N-$NH_4^+$ (mg N $L^{-1}$) | 1.95 | | <d.l. | | 0.010 | |
| TN (mg N $L^{-1}$) | 10.01 | | 0.697 | | 0.862 | |
| TON (mg N $L^{-1}$) | 7.84 | | 0.535 | | 0.625 | |
| SRP (mg P $L^{-1}$) | 0.288 | | <d.l. | | <d.l. | |
| TP (mg P $L^{-1}$) | 0.338 | | <d.l. | | 0.066 | |

**Table S2**. Conditionally rare whole-body bacterial operational taxonomic units (OTUs) associated with *Daphnia* spp. from lakes Koronia and Vegoritida, Greece.

| Phylum | OTUs | % of total OTUs | % OTUs per phylum |
|---|---|---|---|
| Pseudomonadota | 83 | 4.0% | 16.3% |
| Actinomycetota | 49 | 2.4% | 19.3% |
| Bacillota | 37 | 1.8% | 13.4% |
| Bacteroidota | 31 | 1.5% | 11.4% |
| Planctomycetota | 16 | 0.8% | 11.5% |
| Cyanobacteriota | 7 | 0.3% | 9.1% |
| Bacteria unclassified | 6 | 0.3% | 3.2% |
| Verrucomicrobiota | 5 | 0.2% | 6.6% |
| Deinococcota | 4 | 0.2% | 22.2% |
| Abditibacteriota | 2 | 0.1% | 50.0% |
| Campylobacterota | 2 | 0.1% | 50.0% |
| Balneolota | 2 | 0.1% | 28.6% |
| Candidatus Kapabacteria | 2 | 0.1% | 28.6% |
| Thermodesulfobacteriota | 2 | 0.1% | 5.6% |
| Patescibacteria | 2 | 0.1% | 3.8% |
| Fusobacteriota | 1 | 0.0% | 14.3% |
| Chloroflexota | 1 | 0.0% | 2.1% |

**Table S3**. *Daphnia*-associate bacterial operational taxonomic units (OTU) with ≥1% relative abundance and being conditionally rare.

| | ***OTU*** | ***PHYLUM*** | ***CLASS*** | ***ORDER*** | ***FAMILY*** | ***GENUS*** |
|---|---|---|---|---|---|---|
| 01. | 0001 | Bacteroidota | Bacteroidia | Sphingobacteriales | Sphingobacteriaceae | *Mucilaginibacter* |
| 02. | 0015 | Pseudomonadota | Alphaproteobacteria | Rhodobacterales | Paracoccaceae | Paracoccaceae_unclassified |
| 03. | 0034 | Pseudomonadota | Alphaproteobacteria | Pelagibacterales | Clade_III | Clade III insertae sedis genus |
| 04. | 0040 | Bacteroidota | Bacteroidia | Flavobacteriales | Crocinitomicaceae | *Fluviicola* |
| 05. | 0047 | Bacillota | Bacilli | Bacillales | Bacillaceae | *Geobacillus* |
| 06. | 0057 | Planctomycetota | Planctomycetes | Pirellulales | Pirellulaceae | *Pirellula* |
| 07. | 0061 | Pseudomonadota | Gammaproteobacteria | Burkholderiales | Burkholderiaceae | *Polynucleobacter* |
| 08. | 0063 | Cyanobacteriota | Cyanobacteriia | Cyanobacteriales | Nostocaceae | *Dolichospermum* |
| 09. | 0068 | Bacillota | Bacilli | Lactobacillales | Streptococcaceae | *Streptococcus* |
| 10. | 0070 | Cyanobacteriota | Cyanobacteriia | Cyanobacteriales | Microcystaceae | *Microcystis* |
| 11. | 0074 | Pseudomonadota | Alphaproteobacteria | Sphingomonadales | Sphingomonadaceae | *Sphingomonas* |
| 12. | 0086 | Bacteroidota | Bacteroidia | Flavobacteriales | Flavobacteriaceae | *Flavobacterium* |
| 13. | 0087 | Bacteroidota | Bacteroidia | Bacteroidia_unclassified | Bacteroidia_unclassified | Bacteroidia_unclassified |
| 14. | 0091 | Pseudomonadota | Gammaproteobacteria | Burkholderiales | Burkholderiaceae | *Limnobacter* |
| 15. | 0097 | Bacteroidota | Bacteroidia | Flavobacteriales | Flavobacteriaceae | *Flavobacterium* |
| 16. | 0115 | Pseudomonadota | Alphaproteobacteria | Sphingomonadales | Sphingomonadaceae | *Sphingomonas* |
| 17. | 0127 | Bacillota | Bacilli | Lactobacillales | Lactobacillales_unclassified | Lactobacillales_unclassified |
| 18. | 0130 | Pseudomonadota | Gammaproteobacteria | Burkholderiales | Burkholderiaceae | *Ralstonia* |
| 19. | 0138 | Pseudomonadota | Gammaproteobacteria | Enterobacterales | Vibrionaceae | *Vibrio* |
| 20. | 0154 | Deinococcota | Deinococci | Deinococcales | Trueperaceae | *Truepera* |
| 21. | 0157 | Pseudomonadota | Gammaproteobacteria | Lysobacterales | Lysobacteraceae | *Pseudoxanthomonas* |
| 22. | 0172 | Bacillota | Bacilli | RF39 | RF39_insertae_sedis_family | RF39 insertae sedis genus |
| 23. | 0175 | Pseudomonadota | Gammaproteobacteria | Pseudomonadales | Moraxellaceae | *Psychrobacter* |
| 24. | 0179 | Abditibacteriota | Abditibacteriia | Abditibacteriales | Abditibacteriaceae | *Abditibacterium* |

| | | | | | | |
|---|---|---|---|---|---|---|
| 25. | 0190 | Bacteroidota | Bacteroidia | Flavobacteriales | Flavobacteriaceae | *Flavobacterium* |
| 26. | 0196 | Cyanobacteriota | Cyanobacteriia | Cyanobacteriales | Cyanobacteriales_unclassified | Cyanobacteriales unclass. |
| 27. | 0198 | Bacillota | Bacilli | Lactobacillales | Lactobacillales_unclassified | Lactobacillales unclass. |
| 28. | 0208 | Actinomycetota | Actinobacteria | Frankiales | Sporichthyaceae | hgcI clade |
| 29. | 0247 | Patescibacteria | Patescibacteria_unclassified | Patescibacteria_unclassified | Patescibacteria_unclassified | Patescibacteria unclass. |

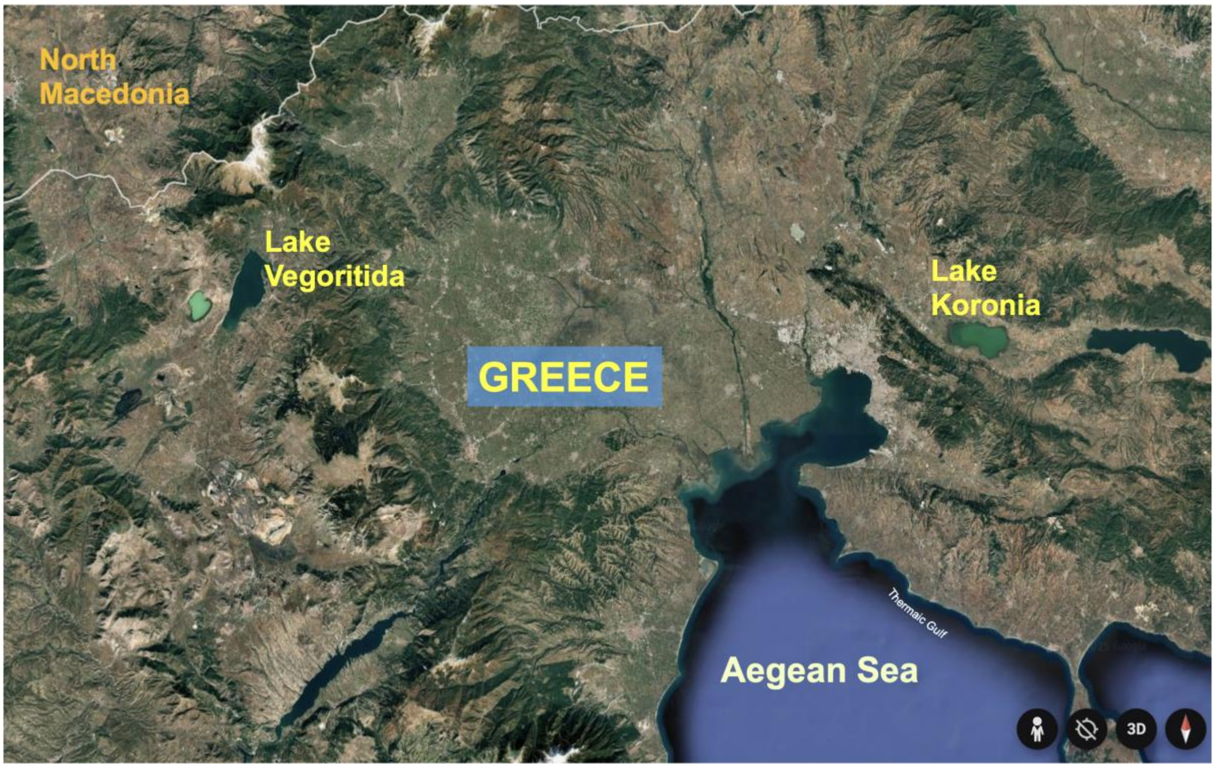


**Figure S1**. Lakes Koronia and Vegoritida.

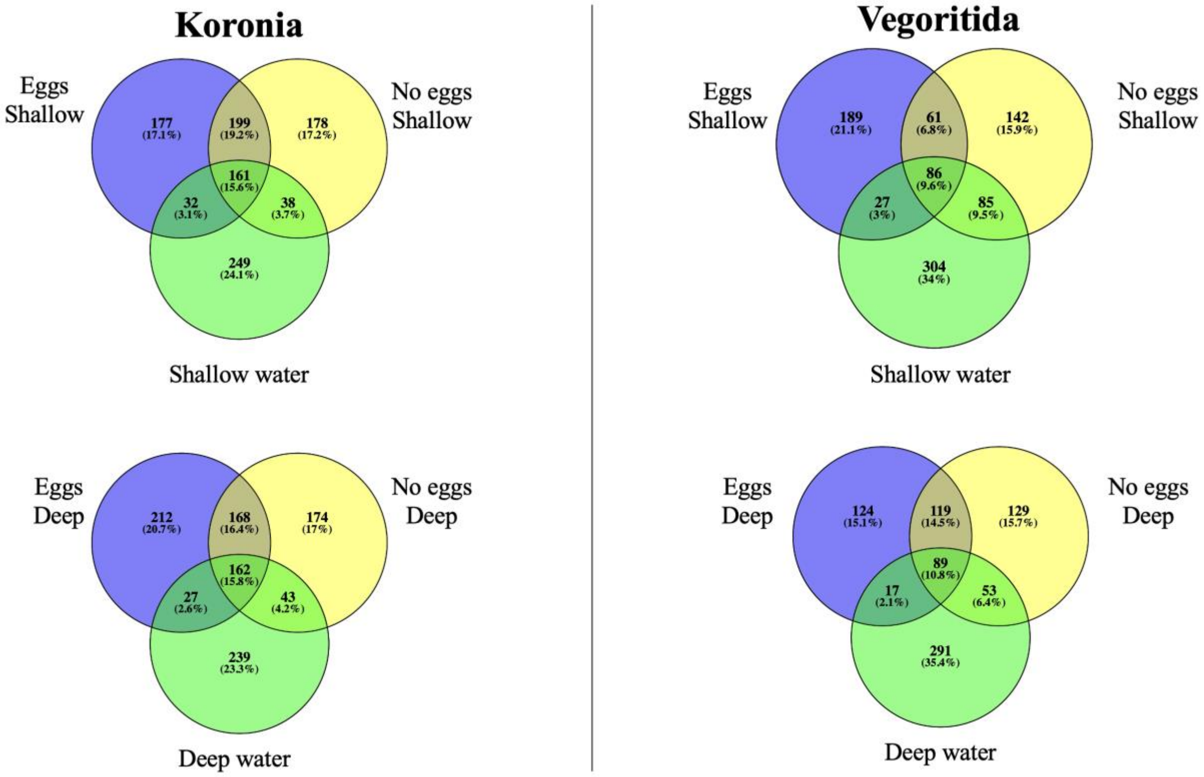


**Figure S2**. Venn diagram of the whole-body bacterial operational taxonomic units (OTUs) associated with *Daphnia* spp. from lakes Koronia and Vegoritida, Greece.
